# Sex-specific Stress and Behavioural Responses to Human Experimenters in Rats

**DOI:** 10.1101/2021.12.01.470834

**Authors:** Jamshid Faraji, Mirela Ambeskovic, Nevyn Sauter, Jaxson Toly, Kera Whitten, Nayara Antunes Lopes, David M. Olson, Gerlinde A. S. Metz

## Abstract

The sex of the experimenter may cause stress in animal models and be a major confounding factor in preclinical research. We studied the effects of the sex of the experimenter on female and male rat anxiety behaviours using thigmotaxis in the open field test, anxiety-induced changes in brain and back temperature using infra-red thermography, and alterations in plasma concentrations of stress hormones, corticosterone and oxytocin. Female rats displayed consistently exacerbated anxiety-related behaviours along with increased infrared cutaneous temperature during repeated exposure to male experimenters. Experimental stress further intensified thermal responses to a male experimenter, especially in female rats. These behavioural responses to a male experimenter in females were associated with higher circulating corticosterone and lower oxytocin levels. Similar responses were induced by a T-shirt worn by a human male. These findings suggest that emotional and physiological responses of female rats to a male experimenter are influenced by visual and olfactory cues. These results emphasize the need to standardize and report experimenter sex throughout a study to avoid ambiguity in interpretation of the results.

## Introduction

Humans 1 and non-human animals 2 are impacted by interpersonal relationships. Laboratory practice with animals typically requires experimenters to build and maintain a close interaction with the subjects, a process that unavoidably involves handling, transferring animals from home cage to the experimental setting and experimental procedures. The same aspects of interaction and manipulation across the experimental course in interplay with the animals’ inherent factors (e.g., age, sex, and species) and environmental influences (e.g., stress, nutrition, rearing and housing conditions) may create an extensive inconsistency in experimental results.

Even when taking particular care about standardizing procedures, controlling for age, sex, litter effects, time of day and season, experimental research still creates situations where inter- and intra-rater concordance and experimenter-dependent manipulations affect the results 3,4. Reasons for this failure include variations in laboratory procedures, the complex nature of research with life rodents, and transgenerational inheritance of ancestral experiences and epigenetic regulators of phenotypic traits 5. In particular, extensive research has documented that stress inevitably confounds measurements and interpretation of experimental results through behavioural, physiologic and metabolic changes linked to elevated activity of the hypothalamic-pituitary-adrenal (HPA) axis 6–8 that may even impact subsequent generations 9–11. Stress is the most salient influence on experimental outcomes, affecting emotion, cognition, movement and sensory function, and underlying inflammatory and physiological response of all organ systems, including the brain 12. In general, female rodents seem to be more vulnerable than males to social and behavioural stressors in terms of circulating oxytocin (OT) responses 13.

It was shown that rodents respond to male but not female human experimenters with a robust stress response and stress-induced analgesia 14. In particular, mice of both sexes observed by male experimenters displayed lower levels of pain than those observed by females 14. The purpose of the present study was to explore sex-specific behavioural and physiological stress responses in a rat model. The results show that female rats are more susceptible than males to the male observer effect. We show that olfactory and visual cues by a male experimenter activates the HPA axis with potentially wide-ranging and lasting effects on behavioural and physiological outcomes. The findings confirm earlier reports 13 of females being more vulnerable than males in terms of endogenous OT secretion in response to social stimuli.

## Results

### Baseline: *Male sex of the experimenter increases anxiety-related behaviours in male and female rats*

Females (n=6) responded to the presence of the male experimenter by more pronounced thigmotactic behaviours and higher variations in the surface thermal outcomes than male rats (n=6). Fig. 1**A** illustrates the thigmotaxis area in the open field which comprised the marginal part of the open zone around the wall. Thigmotaxis in females significantly increased in Experiment 1 when they were required to explore the open field in the presence of a male experimenter (219.34±5.49 vs. 154.33±5.49 s; *F*_1,10_=69.97, *p*<0.001, η^2^=0.87; *Repeated-Measure* ANOVA). We then tested whether this effect could be replicated with T-shirts worn by men in Experiment 2, and in Experiments 3 and 4 in which animals were exposed to the same-sex experimenters and their worn T-shirts. Thigmotaxis in Experiments 2-4, however, revealed no significant differences between females and males (all *p*≥0.05; Fig. 1**B**-**D**). Hence, mainly the emotional aspects of exploration were impacted by the presence of the opposite-sex experimenter only in females. Fig. 1**E** shows thermographic images of two regions of interest [head (left and right) and back] in assessments of surface temperature during open-field exploration. Females and males (n=6/group) displayed different patterns of changes in cutaneous temperatures across all five time bins. Also, the heat change pattern in the ROIs appeared to follow an order of head > back where the head in both groups emitted more heat (females, 33.67±0.27 °C vs. males, 32.78±0.27 °C) than back (females, 31.34±0.09 °C vs. males, 30.51±0.09 °C).

**Figure 1.**
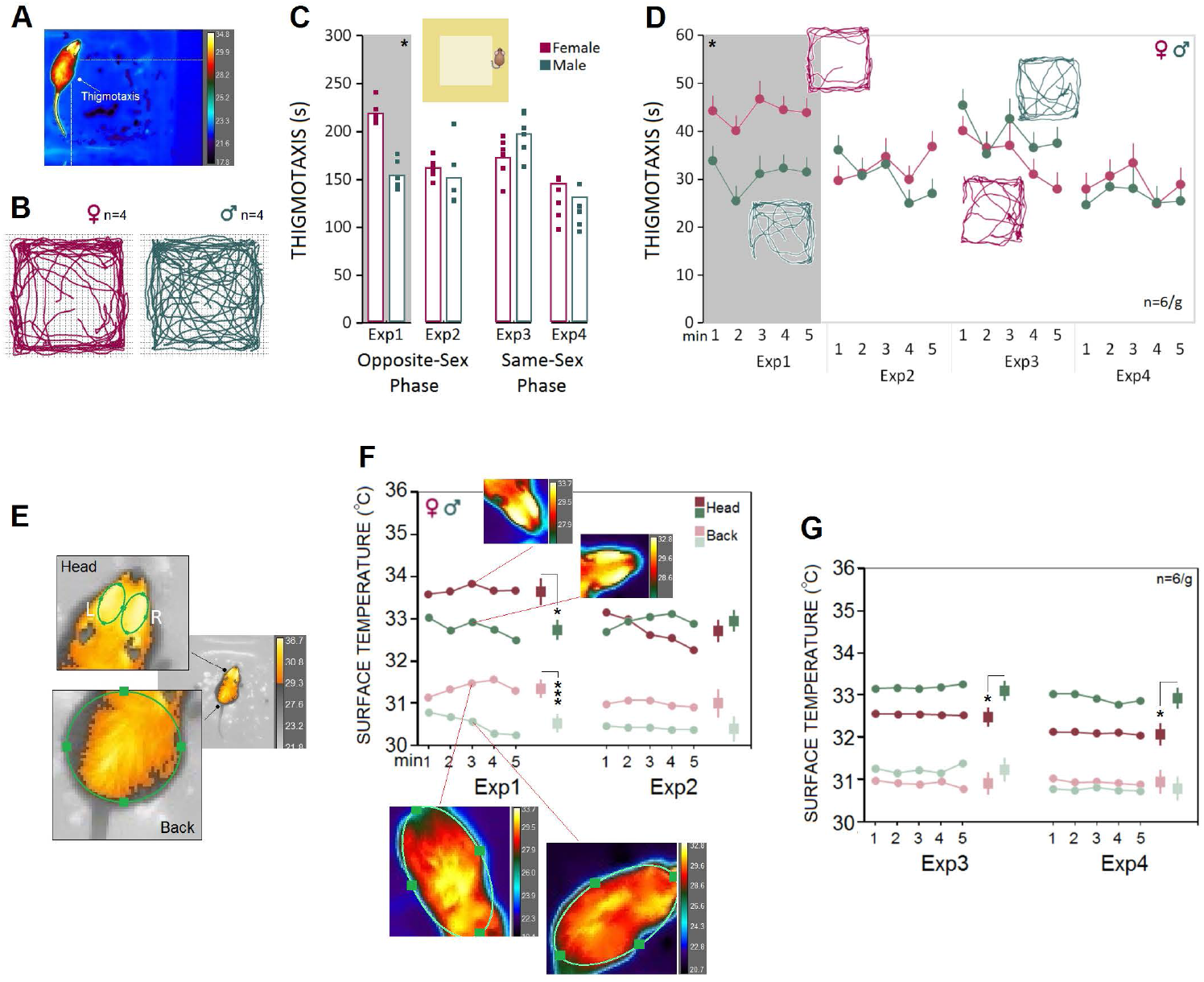
Experimental design. Male and female adult rats were exposed to male and female experimenters in four distinct experiments (Exp1-4). In Experiments 1-2, rats were exposed to opposite-sex experimenters or their T-shirts, whereas rats in the Experiments 3-4 were required to be handled and/or tested by same-sex experimenters. In all experiments animals were manipulated for handling (**A**), pre-stress open field testing (**B**), stress (**C**), stress and post-stress open field testing (**D**), and blood sampling (**E**). Rats’ thermal responses to the experimental procedures were recorded by an infrared (IR) thermographic camera in the presence or absence of the experimenters.

Results showing the thermal changes during open field exploration in Experiments 1 and 2 are displayed in Fig. 1**F**. The heat emitted from head and back ROIs presented a robust effect of Sex (Group) when animals were required to explore the open field in the presence of the opposite-sex experimenters in Experiment 1. Similar to thigmotaxis, compared to males the cutaneous thermal changes in females revealed significantly higher temperatures in the head (*F*_1,10_=8.05, *p*<0.05, η^2^=0.44; *Repeated-Measure* ANOVA) and back (*F*_1,10_=39.82, *p*<0.001, η^2^=0.79; *Repeated-Measure* ANOVA). No effects of Side (left vs right) and Time Bin were found. Also, there was no sex difference in thermal responses to the T-shirts worn by the opposite-sex experimenter in Experiment 2. We thus failed to replicate the sex-specific effect of the male observer on the cutaneous thermal levels in Experiment 2 when animals were only exposed to the T-shirts worn by the opposite-sex experimenters.

However, the profile of thermal changes in Experiments 3 and 4 was different among females and males when animals were exposed to the same-sex experimenters and their T-shirts. In both experiments, male rats displayed an exacerbated thermal response in the head only to the male experimenter (Exp. 3; males, 33.15±0.18 °C vs. females, 32.51±0.18 °C) and the experimenter’s T-shirt (Exp. 4; males, 32.91±0.23 °C vs females, 32.08±0.24 °C) relative to females. MANOVA conducted for the head did show significant effects of Sex (Exp. 3: *F*_1,10_=6.15, *p*<0.05, η^2^=0.38; Exp. 4: *F*_1,10_=6.07, *p*<0.05, η^2^=0.37), but Side (left vs right), Back, and Time Bin (all *p*≥0.05, Fig. 1**G**). Thus, we confirmed that both female and male rats were vulnerable to a male experimenter. Females, however, displayed more exaggerated responses to the presence of the opposite-sex experimenter as shown by the increased thigmotactic behaviours and the surface temperature in the head and back.

### Baseline: *Male rats are more susceptible than females to the presence of a male experimenter during a stressful experience*

Experimental procedures typically impose mild to severe levels of unintended stress upon rodents in laboratory practice and may confound the outcomes of the manipulations. Here we examined how rats that previously experienced stress respond to human experimenters of the same or opposite sex. We chose a single-session psychological stress procedure in the presence of the opposite- and same-sex experimenters and their T-shirts for seven min (day 12). A comparable pattern of thermal changes was found in females and males in Experiments 1 and 2 when animals did experience stress in the presence of the opposite-sex experimenters or their T-shirts (Fig.3**A**) No significant effect of Sex and Time Bin was observed (all *p*≥0.05, Fig. 3**A**, left panel). However, thermal activity in response to the same-sex experimenter and their T-shirts in Experiments 3 and 4 revealed a male-specific response to the male experimenter. Male rats showed higher thermal responses than females in both head (33.87±0.28 C vs. 32.87±0.28 °C; *F*_1,10_=5.94, *p*<0.05, η^2^=0.37; ANOVA) and back (32.09±0.16 °C vs. 31.39±0.16 °C; *F*_1,10_=8.57, *p*<0.01, η^2^=0.46; ANOVA) when stressed in the presence of a female experimenter. When exposed to the T-shirts worn by the same-sex experimenter, however, only heat emitted from back in males indicated a significant difference from females (31.93±0.22 °C vs. 30.88±0.22 °C; *F*_1,10_=10.89, *p*<0.01, η^2^=0.52; ANOVA, Fig. 2**A**, right panel). Thus, it appears that male rats experience greater IR thermal changes than females when exposed to a male experimenter during stress.

**Figure 2.**
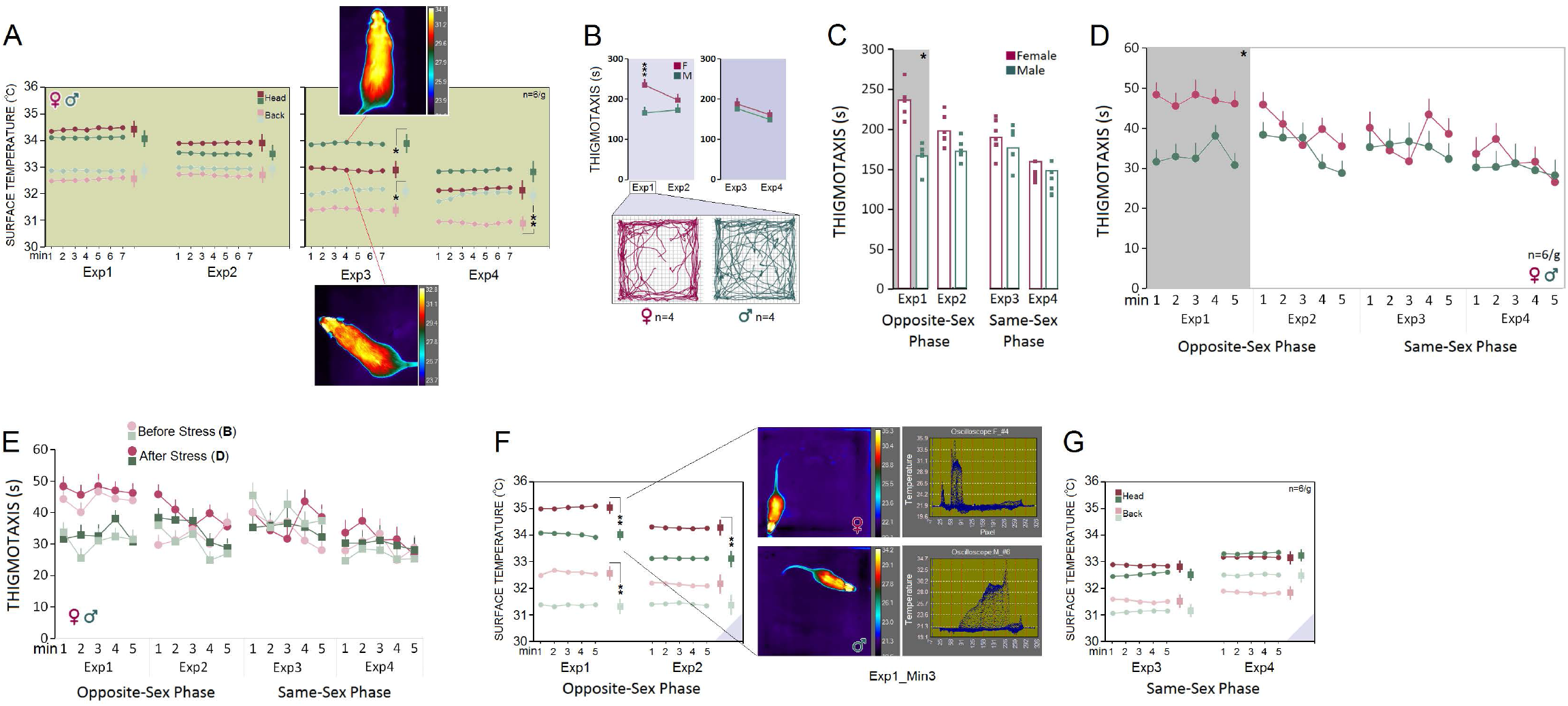
Open field testing and IR imaging. (**A**) Exploratory movements of male and female rats were analysed for thigmotaxis or the repetitive pattern of exploration near to the wall in the open field. (**B-D**) Thigmotaxis in Experiment 1 was significantly impacted when female rats were required to explore the open field in the presence of an opposite-sex experimenter. The multipath shown in the panel B motion track graphics compares thigmotaxis taken from four representative rats during Exp1. Small squares represent individual rats in each group and experimental session. Gray boxes demark statistically significant differences between sexes. (**E**) The IR thermographic imaging showing two regions of interest [head (left and right) and back] in assessments of surface temperature during the open field exploration. (**F**&**G**) Because there were no differences in the thermal responses between the left and right sides of the head, the average of cutaneous temperatures for the left and right sides were used. The heat emitted from both ROIs presented a robust effect of Sex in Experiment 1, as females showed consistently higher cutaneous thermal temperatures than males for the head and back. The inset IR graphic output provide samples of thermal differences in females and males in both ROIs. However, when animals were tested by same-sex experimenters in Experiments 3 and 4, male rats showed higher thermal responses than females in the head. Small squares represent individual rats in each group and experimental session. \**p* ≤ 0.05, \*\*\**p* ≤ 0.001; *one-way* and *repeated-measure* ANOVA, n = 6/group. Error bars show ± SEM.

**Figure 3.**
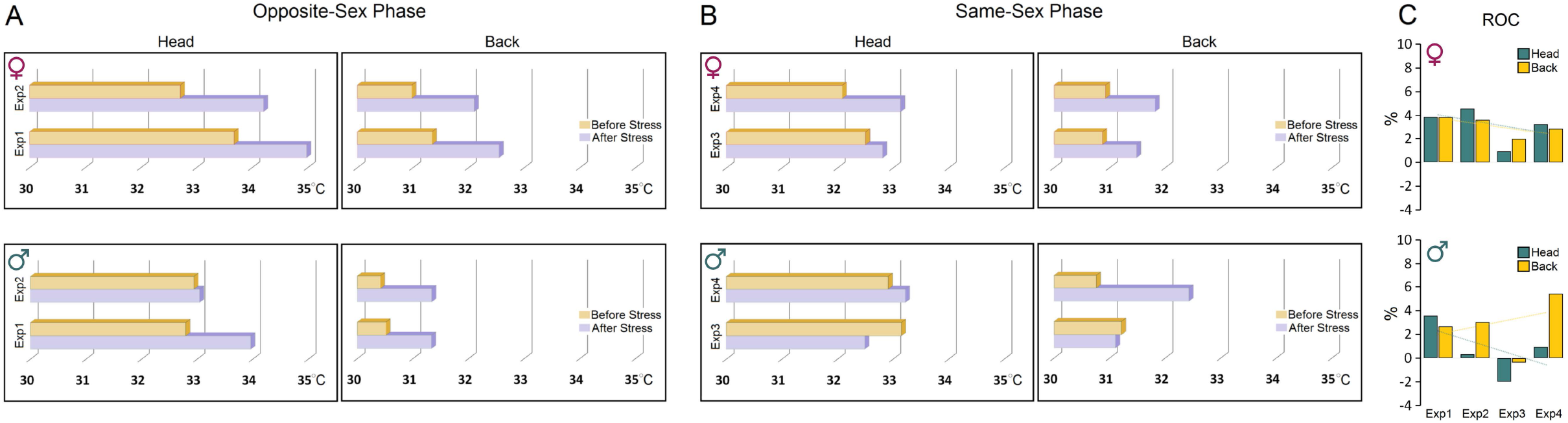
Stress and post-stress open field exploration. (**A**) No significant difference was observed between females and males in the surface temperature when rats were stressed in the presence of opposite-sex experimenters or their T-shirts (Exp1&2). However, thermal responses in Experiment 3 indicated higher cutaneous temperature in males than females when stressed in the presence of same-sex experimenters (right panel; Exp3). Inset thermal graphics compare changes in cutaneous temperature in two representative rats in min 4. Stress in the presence of the T-shirt of the same-sex experimenter also significantly increased the surface temperature of the back in males (Exp4). (**B-D**) Although thermal responses during stress in Experiment 1 showed no sex differences, poststress thigmotaxis noticeably increased in females. The multipath taken from four representative rats in each group is shown in the inset motion track graphics. Gray boxes demark the statistically significant differences between sexes in Experiment 1. (**E**) A comparison of the pre- and poststress thigmotaxis indicated females spent more times near to the wall than males at both time points in the presence of a male experimenter. (**F**&**G**) Interestingly, females displayed higher thermal changes than males in the head and back when exposed to a male experimenter and the male’s T-shirt (Exp1&2) after stress. The middle panel represents thermal changes in min 3 in a female and male rat accompanied by the pertinent oscilloscopes. There were no sex differences in the post-stress same-sex exposure (Exp3&4). Purple triangles in the panels F & G represent *D. IR recording (stress-open field). *p* ≤ 0.05, \*\**p* ≤ 0.01, \*\*\**p* ≤ 0.001; one-way and repeated-measure ANOVA, n = 6/group. Error bars show ± SEM.

### Stress exerts sex-specific effects on emotionality, and intensifies thermal responses to a male experimenter, especially in female rats

To assess the interactions between the stress paradigm and the presence of an experimenter, animals underwent stress in the absence of an experimenter. Immediately after the stress, animals were tested and filmed for IR thermography in the open field in the presence of an experimenter. Thigmotaxis influenced by a 5-min session of elevated platform stress is illustrated in Fig. 2**B-D**. Again, thigmotaxis in Experiment 1 revealed that females spent significantly more time in the thigmotaxis area close to the wall than males in the presence of an opposite-sex experimenter (235.46±7.31 vs. 165.85±7.31 s; *F*_1,10_=45.28, *p*<0.001, η^2^=0.81; ANOVA). No further significant effect of Sex and Time bin was observed in Experiments 2-4 (all *p*≥0.05) where animals were exposed to the T-shirts worn by the opposite-sex or the same-sex experimenters. Also, a comparison between pre- and post-stress thigmotaxis showed no differences between females and males (all *p*≥0.05, Fig. 2**E**) indicating that the transient stress paradigm did not change emotionality in the open field.

Similar to the baseline thermal responses, however, female rats responded to the male experimenter and the male-worn T-shirt with an exacerbated cutaneous thermal reaction when compared with male rats after stress (Fig. 2**F**&**G**). The heat emitted from head (34.98±0.18 °C vs. 33.95±0.18 °C; *F*_1,10_=14.87, *p*<0.01, η^2^=0.59; ANOVA) and back (32.55±0.26 °C vs. 31.33±0.26 °C; *F*_1,10_=10.59, *p*<0.001, η^2^=0.51; ANOVA) presented a robust effect of Group in Experiment 1 where the cutaneous thermal responses to the male experimenter was significantly more exaggerated in females than males. Thermal changes in Experiment 2 also were higher in females than males in the head (34.18±0.25 °C vs. 33.04±0.25 °C; *F*_1,10_=10.44, *p*<0.01, η^2^=0.51; ANOVA) in response to the T-shirt worn by male experimenters. The significant effect of Group, however, disappeared in Experiments 3 and 4 when animals were required to explore the open field in the presence of the same-sex experimenter and the experimenters’ T-shirts (all *p*≥0.05). No effect of Time bin was observed across all experiments (all *p*≥0.05).

Further comparisons of thermal changes prior to and after stress showed that both groups were susceptible to the stress procedure when responding to the opposite-sex experimenters (Fig. 3**A**&**B**). However, the profile of thermal changes before and after stress in both ROIs in females was noticeably different from males indicating higher vulnerability of female thermal responses to the experimenter’s sex. Females’ susceptibility to the transient stress and the experimenter sex was also supported by additional analysis of the rate of changes (ROC) showing that female rats experienced larger changes in cutaneous temperatures after stress in response to experimenter sex (Female [*head*]: 12.65% vs. Male [*head*]: 2.83%; Female [*back*]: 12.31% vs. Male [*back*]: 10.78%, Fig.3**C**). Thus, although behavioural responses to the experimenter sex remained unchanged after stress, the cutaneous thermal responses to both, opposite- and same-sex experimenters in females were noticeably affected by the transient stress.

### CORT and OT: *Presence of male experimenters or their T-shirts raised plasma CORT levels and reduced OT levels in female rats*

The male experimenters and male-worn T-shirts induced sex-specific behavioural and neurophysiological changes that can be linked to higher anxiety-like behaviours and stress responses in rats. Here, we examined stress-induced activation of the HPA axis in circulating plasma CORT changes linked to experimenter sex. One female and one male rat were excluded from the CORT analysis due to technical issues. Generally, females had higher plasma CORT levels than males in the presence of a male experimenter (Mean Rank: 8.83 vs 4.17; *U_6,6_=4.000, Z*=-2.24, *p*<0.05; Mann-Whitney *U*) or their worn T-shirts (Mean Rank: 8.67 vs 4.33; *U_6,6_=5.000, Z*=-2.08, *p*<0.05; Mann-Whitney *U*) across the four experiments. The between-group differences disappeared in Experiment 3 when animals were exposed to a same-sex experimenter (Mean Rank: 6.00 vs 6.00; *U_5,6_=15.000, Z*=0.00, *p*=1.000; Mann-Whitney *U*) or their worn T-shirts (Mean Rank: 7.92 vs 5.08; *U_6,6_=9.500, Z*=-1.36, *p*=0.17; Mann-Whitney *U*), although CORT values in females were still higher than those in males (Fig. 4**A**). Overall, the HPA axis response to the experimenter was limited to female rats only in the presence of male experimenter or T-shirts worn by men.

**Figure 4.**
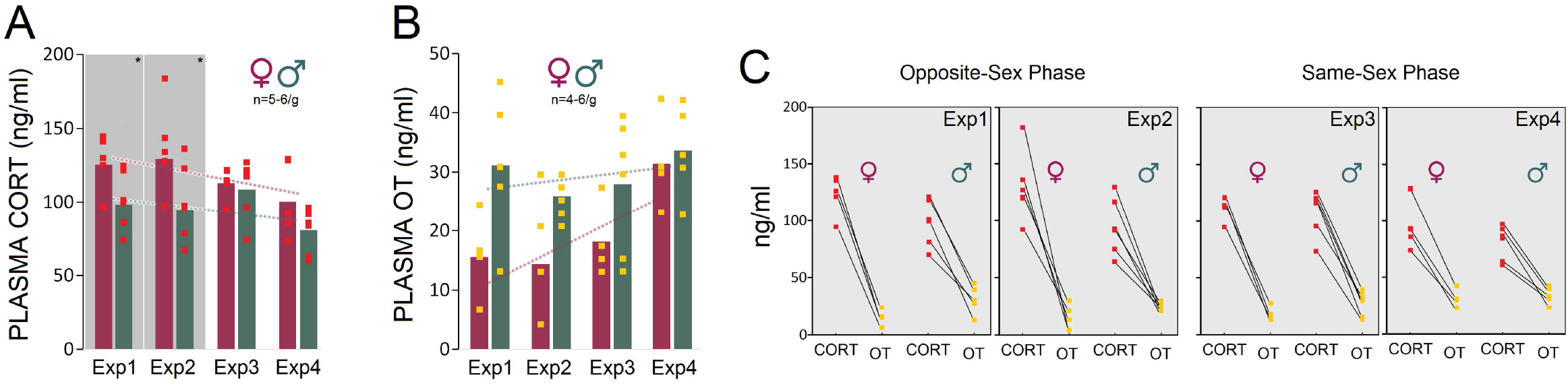
(**A**&**B**) *Thermal changes prior to and after stress*. Both groups were susceptible to the stress procedure in the presence of opposite-sex experimenters. However, female thermal responses in both ROIs were noticeably different from males before and after stress indicating higher vulnerability of females to the impact of stress and experimenter sex. (**C**) The rate of changes (ROC) provides further support for exacerbated thermal responses in females to experimenter sex after stress. Note the trendlines of thermal changes in the head and back in both sexes that noticeably depict sex-specific differences in the regional thermal responses to stress and experimenter sex.

We also hypothesized that elevated CORT induced by a male experimenter may dampen oxytocinergic influences that are linked to positive social interactions. The presence of a male experimenter moderately reduced the plasma OT levels in females (15.53 ng/ml) compared to males (31.15 ng/ml) in Experiment 1 (Mean Rank: 3.25 vs 6.40; *U_4,5_=3.000, Z*=-1.71, *p*=0.08; Mann-Whitney *U*) and experiment 2 (14.37 vs 25.93 ng/ml, Mean Rank: 4.00 vs 7.67; *U_5,6_=5.000, Z*=-1.84, *p*=0.06; Mann-Whitney *U*) when female rats were exposed to men-worn T-shirts. Despite the decline in female OT levels in the presence of a male experimenter, the consistently higher OT levels in male rats across experiments indicated that the experimenter sex only had a marginal impact on the males’ oxytocinergic responses (Fig. 4**B**&**C**).

### Correlational analysis: *Behavioural responses to stress and the experimenter sex do not predict thermal changes*

Because there were no hemispheric differences, the average of cutaneous temperatures for the left and right sides of the head were used for correlational analyses. Pearson’s correlation analysis revealed no significant relationship between thigmotactic behaviours in the open field and changes in surface temperature in the corresponding ROIs *before stress* (Fig. 5**A**), except for the back when females were exposed to a male experimenter (Exp.1, *r*= −0.93, *p*=0.01). Also, the only significant correlation between thigmotaxis and thermal responses *after stress* was observed in Experiment 1 for females in the head (Exp.1, *r*= −0.90, *p*<0.01; Fig. 5**B**). Thus, in most cases changes in thigmotaxis in the open field, either before or after stress, did not predict surface temperature in response to opposite- and the same-sex experimenters. Fig. 5**C** also illustrates correlations between the heat emitted *during stres*s (*C-Baseline*) and *post-stress* exploration in the open field (*D*). Changes in surface temperatures during stress only predicted surface temperatures for the back in males when exposed to a T-shirt worn by a female (Exp.2, *r*=0.87, *p*<0.05) and to a male experimenter (Exp.3, *r*= −0.87, *p*<0.05). In females, however, the correlations between the pre- and post-stress cutaneous temperatures only were significant in the head when they were exposed to T-shirts worn by female experimenters (Exp.4, *r*=0.89, *p*<0.01) suggesting that stress-induced thermal changes did not reliably predict post-stress thermal responses to the experimenter sex in both groups.

**Figure 5.**
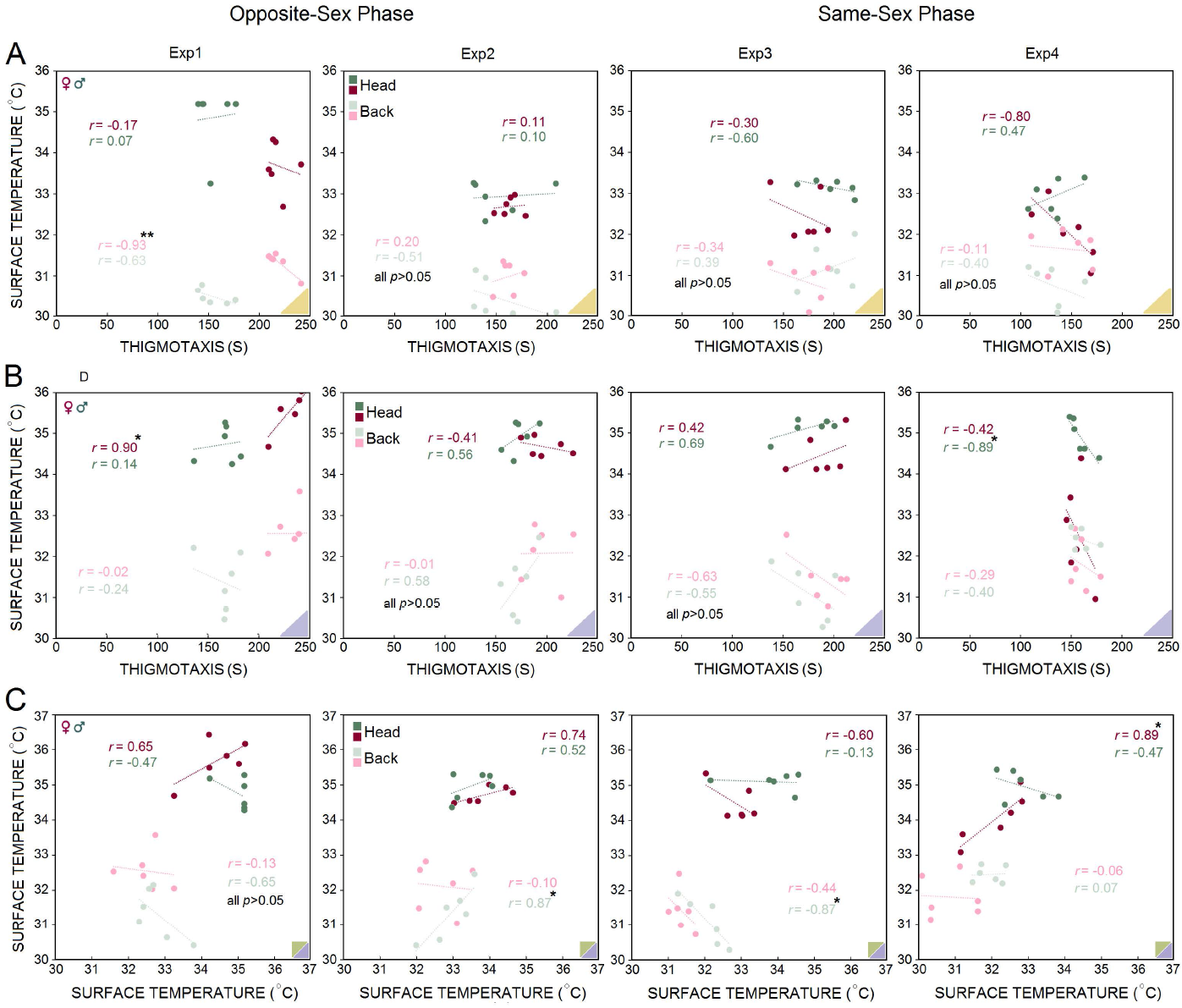
Corticosterone and oxytocin responses to stress and experimenter sex. (**A**) Average CORT levels in females across all experiments were higher than males. Exposure to an oppositesex experimenter significantly elevated plasma CORT levels in female rats only. There were no significant differences between females and males in Experiments 3 and 4 when exposed to samesex experimenters or their T-shirts. Red squares represent individual animals in each group. Gray boxes represent statistically significant differences between sexes (*p*≤0.05, Mann-Whitney *U*; n = 5-6/group) (**B**) Female rats responded to the presence of a male experimenter with reduced plasma OT levels. In contrast, male rats experienced marginal changes in plasma OT levels across experiments (n = 4-6/group). (**C**) Ladder plots (pairwise changes) illustrate that increased CORT levels are associated with reduced OT levels in both sexes, although female rats experienced higher levels of the CORT and lower levels of OT than males particularly in the presence of human male experimenters or their T-shirts.

Moreover, Spearman’s rho correlation coefficient showed that increased plasma CORT can reliably predict increased thigmotactic behaviour indicating that the experimenter sex mediates both the HPA-axis activity and anxiety-like behaviour, particularly in female rats (Fig. 6**A**). The observed correlation in females however was stronger than male rats in Experiments 2 (*r*_s_=1.000, *p*<0.001 vs. *r*_s_=.42, *p*=0.397, n=4-6/g) and 3 (*r*_s_=1.000, *p*<0.001 vs. *r*_s_=.94, *p*<0.01, n=4-6/group). In contrast, there was no correlation between CORT and surface temperature across experiments suggesting that in both sexes there are likely two distinct neurohormonal pathways that may determine the thermal and the HPA-related responses to the experimenter sex (Fig.6**B**). Further analysis also revealed a significant negative correlation between CORT and OT levels exclusively in females in Experiment 1 (*r*_s_= −1.000, *p*<0.01, n=4; Spearman’s rho) and 2 (*r*_s_= −1.000, *p*<0.01, n=5; Spearman’s rho) where increased CORT levels were associated with reduced OT levels in the presence of a male experimenter or male-worn T-shirts. No significant correlations were found between CORT and OT levels in Experiments 3 (*r*_s_= −.667, *p*=0.22, n=5; Spearman’s rho) and 4 (*r*_s_= −.800, *p*=0.20, n=4; Spearman’s rho) in females (Fig. 6**C**).

**Figure 6.**
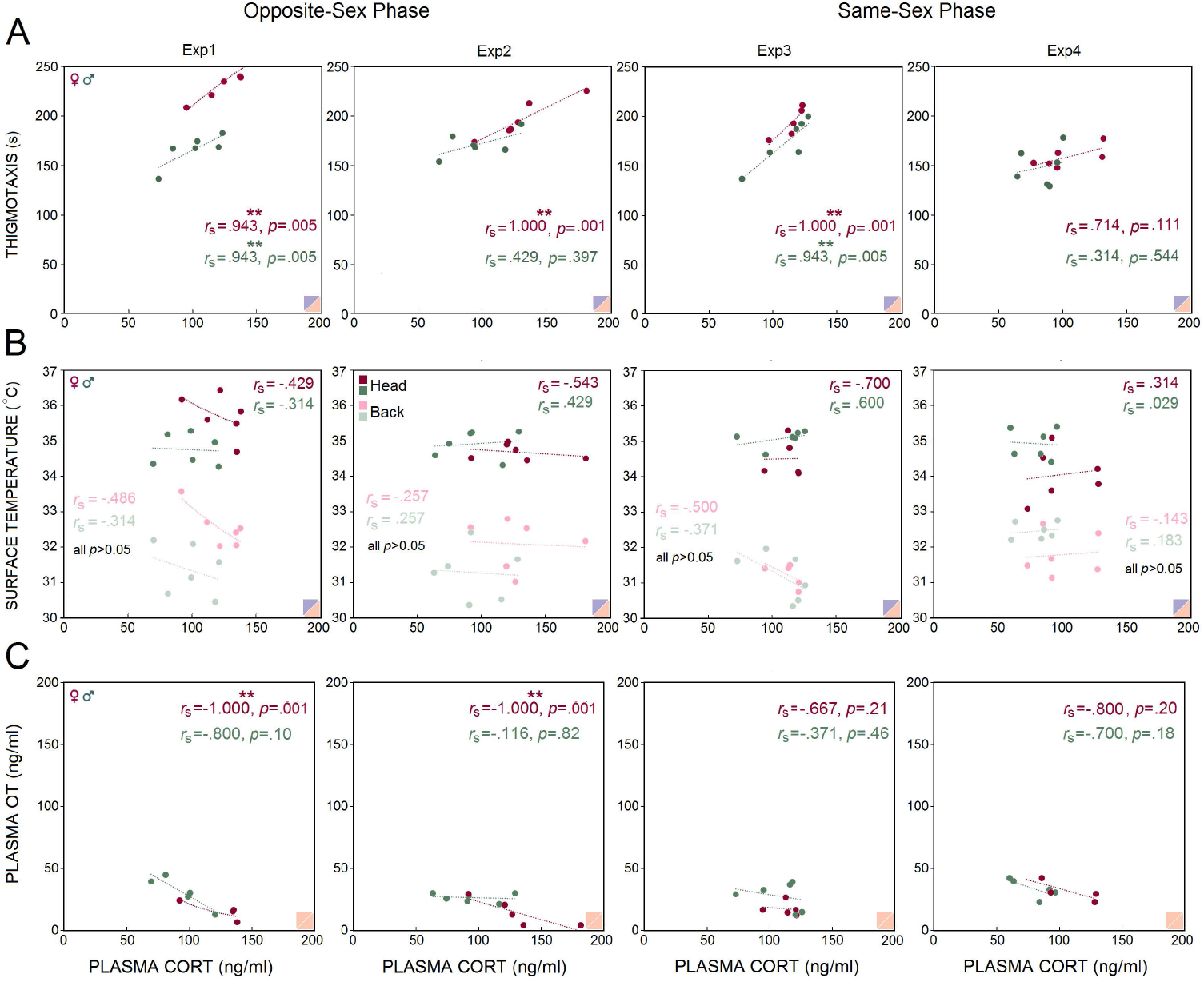
Correlational analysis. Most trials in Experiment 1 to 4 revealed no significant correlation between surface temperature and thigmotaxis before and after stress (panels **A**&**B**). Stress-induced thermal changes in general did not predict post-stress thermal responses to the experimenter sex in both groups (panel **C**). Yellow, green and purple triangles in the corners represent *B. Baseline, C. Baseline*, and *D. IR recording* (*stress-open field*), respectively. \**p* ≤ 0.05, \*\**p* ≤ 0.01; Pearson’s correlation, n = 5-6/group.

## Materials and Methods

### Animals

Male and female Long Evans rats, 8-10 weeks old at the beginning of the experiment, were used in this study. The animals were housed in trios of the same sex in standard housing under a 12:12 h light/dark cycle with light starting at 07:30 h, and food and water provided *ad libitum*. The room temperature was set at 22 °C, and experimental procedures were conducted during the light phase of the cycle at the same time of day. All animals were habituated in their assigned housing condition for two weeks before any experimental manipulation commenced. Aspen wood chips mixed with shredded paper bedding material was used in all home cages and changed once per week by a female animal care staff. Animal care personnel were recommended to minimize their physical contacts with animals while providing animal husbandry. Animals were briefly removed from their cages for 2-3 min when changing the bedding material was necessary. All procedures were approved by the University of Lethbridge Animal Care Committee in compliance with the standards set out by the Canadian Council for Animal Care (CCAC).

### Experimental Design

Figure **7** (**A-E**) illustrates the time course of experimental manipulations in four experiments. *Opposite-sex phase* (*Experiment 1*): The female rats (n=6) were handled and tested by a male experimenter. The male rats (n=6) were handled and tested by a female experimenter. Rats were handled daily for a total of ten days (Days1-10). Each rat was also handled for 2 min prior to any experimental manipulations on the four subsequent days. The experimenter remained in the room during testing, quietly positioned approximately 50 cm from the animal and testing apparatus (Days 11-13). Animals were handled by the same experimenters for two minutes per rat before being anaesthetized with isoflurane for blood sampling (Day 14). *Opposite-sex phase* (*Experiment 2*): Experiment 2 was carried out in the same manner as of Experiment 1, but instead of the experimenter being present in the room, a cotton T-shirt was placed on a chair approximately 50 cm away from the animal 14. The T-shirt was worn by the experimenter for at least 12 h prior to the assessments. The *Same-sex phase* (*Experiment 3*): All procedures were identical to Experiments 1 and 2, except that female rats (n=6) were handled and tested by a female experimenter and male rats (n=6) were handled and tested by a male experimenter. *Same-sex phase* (*Experiment 4*): All procedures were identical to Experiment 3 except that animals were tested in the presence of the experimenters’ T-shirt while the experimenters remained outside of the room for each assessment. Again, the T-shirts were previously worn by the experimenter for at least 12 h beforehand. For all testing procedures, rats were transported in their home cages to a designated test room. Male and female rats were tested and filmed in separate rooms.

**Figure 7.**
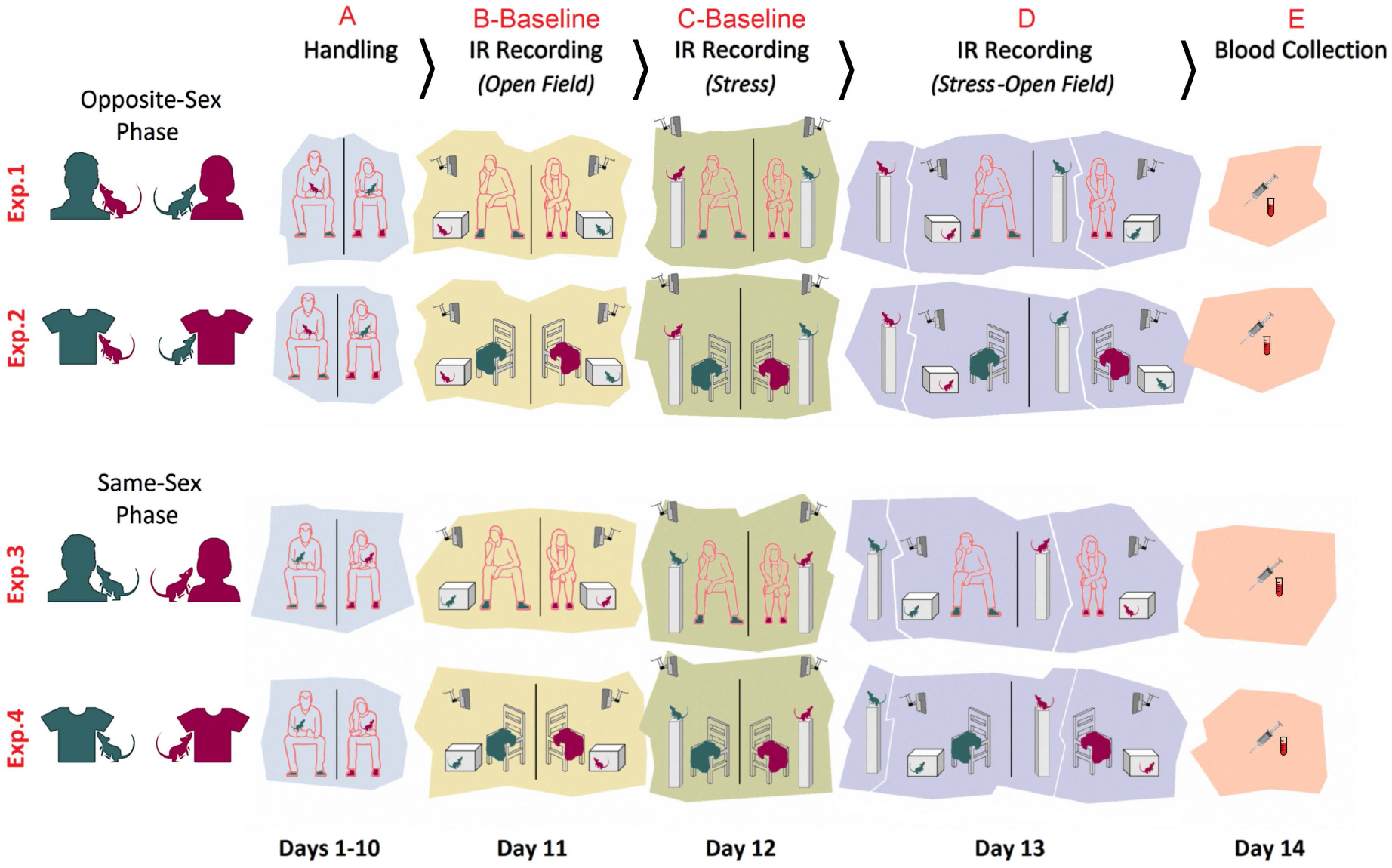
Correlational analysis. (**A**) Plasma CORT and thigmotaxis were positively correlated in both sexes, particularly in female rats. (**B**) There was no significant correlation between the cutaneous temperature and CORT. (**C**) A significant negative correlation was found between the plasma CORT and OT in female rats in Experiments 1 and 2, whereas the correlation between the CORT and OT in male rats was not significant. Purple and pink triangles in the corners represent *D. Stress* and *E. Blood Collection*, respectively. \*\**p* ≤ 0.01; Spearman correlation, n = 4-6/group.

### Handling

In all experiments, animals were handled individually for 5 min daily by their male or female experimenters for a total of 10 days prior to any experimental manipulations. The long-term handling protocol ensured habituation of the rats to the experimenters. The rats were handled in the housing room in a seated position on the lap of the experimenter for two min and then handled near the chest of the experimenter for an additional three min to facilitate olfactory habituation. Experimenters used their same own lab gown during all handling and test sessions.

### Open Field Task

The open field task (OFT) was used to assess anxiety-like behaviour in rats. The apparatus consisted of a square box (50×50 cm) made of transparent Plexiglas and surrounded by walls (45 cm height). Each rat was individually placed at the centre of the box and video recorded for 7 min with an infrared (IR) thermographic camera mounted above the open field. The animals’ movements were analysed for thigmotaxis (time spent close to the walls; 8 cm width) by an experimenter blind to experimental conditions. Thigmotactic behaviour was analysed as an indicator of anxiety and emotionality in the open field 7. After testing each animal, the apparatus was cleaned with 1% Virkon (Antec International Ltd., Suffolk, UK).

### Stress Procedure

The elevated platform stress protocol used as a mild emotional stress was modified from that described previously 15. Animals were individually placed on the platform (20 cm × 20 cm on a Plexiglass stand 1 m above ground) for a single 7-min session. After the animal was placed on the platform and while it was filmed, the experimenter was quietly sitting (Exp. 1&3-C), or the experimenter’s T-shirt was placed (Exp. 2&4-C) on a chair a short distance (~50 cm) away from the stand. Animals were removed from the platform and returned to the home cage by the experimenter when the stress session was terminated, and the platform was cleaned with 1% Virkon (Antec International Ltd., Suffolk, UK) between stress sessions.

### Infrared (IR) Thermal Imaging

The pre-stress thermal imaging was performed immediately after a 2-min handling session. A FLIR IR thermographic camera (FLIR T450sc, Sweden; fixed emissivity=0.98 specified for skin in the manufacturer’s emissivity table) mounted on top of a transparent Plexiglas box or an elevated platform recorded the cutaneous temperature in rats. IR imaging occurred in a windowless room with a steady temperature set at 22 °C and a relative humidity of ~50%. Animals were protected from direct ventilation.

As previously reported 6, animals were placed individually at the center of the Plexiglas box in a prone position, and the IR thermographic recording was performed above the box without lid because IR radiations are blocked by Plexiglas or stainless steel. The camera was placed ~80 cm above the animal and was able to follow changes in the animal’s surface temperature and its immediate surrounding with thermal resolution of 320×240 pixels per image, thermal sensitivity of <30 mK at 30 °C, and 60 Hz acquisition rate.

IR thermal profiles were then saved and analysed using the FLIR image processing software (FLIR ResearchIR Max software 4.40.6.24). For the purpose of the thermal analysis, two principal regions of interest (ROI) were chosen. (1) *Head* (including eyes), covering a major portion of the frontal and parietal surfaces. An approximate measure of the sagittal suture allowed the elliptic ROI to split the top of the head up into left and right sides. Also, a small segment of interparietal bone was included in either right or left ROIs. To control the effect of the position of each animal on the emitted thermal irradiations, the best postural condition for the head was chosen when rats were moving with their head oriented straight ahead without deviation to the side. (2) *Back*, an oval-shape ROI included lower thoracic and upper and lower lumbar levels extended to the abdominal parts at equidistance from approximately 2.5 cm off the midline. In total, three ROIs (head [left and right] and back) were considered for analysis of changes in surface temperature. For sampling, 5-7 frames representing 5-7 time bins (one frame per each minute of the IR imaging) adjusted to the corresponding ROIs from the head and back were chosen for each animal. ROI sizes were identical for all frames and rats. The best-fit area to the ROIs in each frame/time bin was determined on the basis of the animal’s dorsal posture among nearly 1,750 single frames when approximately all relevant regions were bounded by the radius of the ellipses and/or when the animal was found in a prone position with all four limbs on the ground.

### Blood Samples, CORT Assessment, and Oxytocin Assay

Blood samples were collected in the morning hours between 9:00 and 11:00 am. Briefly, rats were transported individually to the surgical suite and were handled by their assigned male or female experimenters for 2 min prior to anesthesia with 4% isoflurane. During 2–3 min of anesthesia, 0.5-0.9 ml of blood was collected from the tail vein using a heparinized butterfly catheter. Blood samples were taken by a separate male experimenter while the animals were anaesthetized. Blood was then transferred to centrifuge tubes and plasma was obtained after centrifugation at 5,000 rpm for 5 min. The plasma samples were stored at −80 °C until analyzed for CORT and OT concentrations. Plasma CORT levels were determined by enzyme-linked immunosorbent assay (ELISA) using commercial kits (Cayman Chemical, Ann Arbor, MI, USA). Plasma OT levels were determined by enzyme immunoassay (EIA) using a Human/Mouse/Rat Kit (RayBiotech Inc, Norcross, GA, USA) according to the manufacturer’s protocol. The OT concentration (ng/ml) was determined by plotting the mean absorbance of each unknown sample on the standard curve (range 0.1-1,000 ng/ml). The minimum detectable concentrations of OT was 3.6 ng/ml. The Intra- and inter-assay coefficient of variation (CV) were <10% and <15% respectively, as reported by the manufacturer.

### Statistical Analysis

Effects of main factors (Experiment—four levels; Sex—two levels) were analyzed as independent variables for the thigmotaxis in the open field and the surface temperature in different ROIs and frames as dependent variables by *repeated measure*, *one*- and *multivariate* ANOVAs. *Post-hoc* Tukey test was used to adjust for multiple comparisons when multi-level factors (e.g., experiments and frames) were needed to be compared. Familywise error was considered prior to the multiple *post*-*hoc* analyses, if necessary. Also, to evaluate the magnitudes of the effects of experimental manipulations (here, elevated platform stress and experimenter sex) on thigmotaxis, body surface temperature, effect sizes (*η*^2^ for ANOVA) were calculated. Values of *η*^2^= 0.14, 0.06, and 0.01 were considered for large, medium, and small effects, respectively. Moreover, because the plasma CORT and OT values in the present study were not normally distributed, the Mann-Whitney U, a rank-based nonparametric test was used to compare means of the two groups (females vs. males) for a single dependent variable, either CORT or OT. Correlations between variables (surface temperature, thigmotaxis, CORT and OT) were analyzed by Pearson product-moment and Spearman rank-order correlation coefficients. In all statistical analyses (IBM SPSS statistics, Version 21, USA), a p-value of <0.05 (two-tailed) was chosen as the significance level. Results are presented as mean ± standard error.

## Discussion

The replication crisis represents a significant threat to the advancement of the life sciences and medicine 16. The failure to reproduce results requires urgent attention to identifying the variables that can limit reproducibility of research in animal models. The impact of the sex of the experimenter on behavioural and biological processes in laboratory rodents is poorly understood and generally underestimated. Here we show that male and female rats display robust behavioural and physiological stress responses to the sex of the experimenter. A human male experimenter induces more variations in female rats with changes that are linked to the dysregulation of the HPA axis and psychophysiological distress. These fundamental changes in the stress response may potentially influence the results of basic science and preclinical studies, emphasising the need for consideration in experimental design, reporting of the research, and data interpretation.

The HPA axis represents a neurohormonal hallmark of emotionality in humans and animals 12,17, and its activity is characterized by prominent sex differences. In fact, females initiate the HPA-axis activity more rapidly in response to stressful stimuli and produce a greater output of stress hormones 18–20. It appears that sex differences in animal research are not displayed consistently. A growing body of evidence show that sex differences in rodents stem, at least in part, from organizational effects of sex hormones 21, rearing/housing conditions 22, and experimental protocols and parameters 23. Hence, the present findings shows that the neurohormonal differences in the rats’ response to the experimenter sex that appear most pronounced in an opposite-sex framework. Therefore, animal studies involving male experimenters will more likely suffer from experimental bias.

IR thermal measures provide another sensitive indicator of the stress response, which were also impacted by experimenter sex. This observation reflects the sex-specific nature of thermoregulatory activity in rats 6. Indeed, the peripheral autonomic nervous system that regulates perspiration and surface blood perfusion, predominantly determines heat patterns and gradients during aversive experiences 24. Furthermore, HPA-system activity alone does not fully display a complete picture of sex differences during emotional disturbances as neuroendocrine stress-related responses in female rodents are different than males 19. Thus, cutaneous temperature variations may serve as an alternative physiological marker for stress, fear, tension and anxiety 25–27. Here, females displayed greater levels of IR thermal susceptibility to the ‘male observer’ effect 14 when briefly stressed, even though the stress paradigm used in the current study was not salient enough to experimentally induce a prominent stress response in rats. This supports our earlier findings in mice that (1) thermal responses function partially independently of CORT levels, (2) may be more sensitive to subtle effects of stress, and (3) show greater effects in females 6. The findings suggests a distinct neuroregulatory system in thermal response to emotionally threatening stimuli in rats. The present findings also show that synergy between a stressful stimulus and the presence of a male experimenter may further sway experimental outcomes. The experimenter sex may account for a significant portion of the variance of behavioural and physiological observations particularly in female animals.

In regard to the sex-biased thermal alterations and the OT inhibition in response to experimenter sex two mechanistic possibilities can be hypothesized. First, changes in the peripheral temperature in females that are modulated by central subsystems are sex-specific hormone-dependent responses mainly influenced by estrogens 28,29. In parallel with its thermal consequences, estrogen is also expected to centrally increase OT signaling 30, which eases the anxiolytic oxytocinergic function. This OT-mediated signal typically reduces the inhibition inherent to social encounters. The second possibility, alternatively, points to the inhibitory effect of glucocorticoid action on the OT function 31,32 which is influenced by sex steroids, especially estrogens 18. OT normally enhances the glucocorticoid response to acute aversive experiences 33,34. However, increased circulating CORT levels due to stress may reversely inhibit OT-mediated anxiolytic influences 34,35 depending upon the stress regimen and intensity. For instance, a prolonged stressful experience can reduce the anxiolytic effect of OT. It appears that the up- or down-regulation of OT signaling by HPA axis activity not only depends on the brain region, but also on the duration of the stressor (acute vs. chronic) 34. Hence, the neuroendocrine pathways causing variations in thermal maps and reduced OT levels may contribute to sex disparities in response to the presence of a male experimenter via a close interaction with the HPA axis 35.

The present study showed inhomogeneous stress responses towards T-shirts worn by a male experimenter. A number of possibilities should be considered as the main source of discrepancy between the present results and earlier findings 14. First, the T-shirts worn by males may only mimic the presence of an experimenter when specific neurophysiological measures such as pain behaviours and analgesic changes are addressed. Second, olfactory stimuli (e.g., axillary secretions) may differently affect female and male rats and male-associated olfactory inputs 36 to induce greater alterations in females when combined with visual stimuli. Thus, stress-related emotionality and physiology in female rats may be influenced by olfactory cues of male perspiration 14, by visual cues of a present threat 7 or by other neurohormonal stimuli 37,38.

An important corollary to repeated exposure to a male experimenter is the development of habituation, a normal pattern that indicates gradual coping with the stress-induced homeostatic disruption 39. In contrast, a failure to habituate typically represents greater stress vulnerability. The impact of a male’s presence on experimental outcomes in the current study did not diminish over time, although animals were handled by the same- or opposite-sex experimenters for 10 days before and during the experiments. Accordingly, the disparate responses to male experimenters seen in female and male rats were maintained for the full 14 days of repeated exposure. This observation negates the potential that repeated exposure to a male experimenter prior to an experimental manipulation may induce stress tolerance or habituation 39. However, data in the present study shown by the elevated levels of CORT in females in the first two experiments do support an alternative hypothesis that HPA-axis response, in specific conditions, fails to habituate to continuous stressor exposure 40–42.

## Conclusion

The present study demonstrates that male and female experimenters represent a critical variable in rodent models, with female rats being more susceptible than males to the male observer effect. Moreover, the confounding effect of the male experimenter neither diminishes overtime, nor can it be merely attributed to human olfactory stimuli. We showed that the presence of a male experimenter activates the HPA axis with potentially wide-ranging effects on behavioural and physiological outcomes. The findings confirm earlier reports of females being more vulnerable than males in terms of endogenous OT secretion in response to social stimuli 13. Notably, the opposite-sex dynamics that has been recently reviewed in humans 43, may also be applicable to animals in preclinical studies.

In line with an earlier report 14, the present data are thought to encourage committed efforts to solve the replication crisis in the life sciences. Because experimenter sex in animal studies is not consistently reported in the scientific literature, improved standards should require researchers to report the sex of experimenters. In addition, the present findings emphasize the critical importance of including females in preclinical research to address potential sex differences within a translational research framework 13,44,45.

Since the close interaction between the subject and experimenter impacts experimental outcomes, a hands-off test system that collects data throughout the day and night away from the influence of a human experimenter may have the capacity to minimize the experimenter sex effect 46. Thus, the idea of a virtual experimenter 43,47,48 that conceptualizes the application of an automated protocol and computer-based treatment in experiments, may pave the way to standardize experimental procedures and reduce artifacts in preclinical findings.

## Acknowledgements

The authors acknowledge support by the Natural Sciences and Engineering Research Council (NSERC) of Canada Discovery Grants #5628 and #31 to GM, and by the Canadian Institutes of Health Research (CIHR) Project Grant #165890 to DMO.

## Author contributions

JF, and GASM designed the study. JF, MA, NS, JT, KW, NAL performed the experiments and analyzed the data. JF wrote the paper. MA, NAL, DMO and GASM edited the paper. DMO and GASM acquired the funding and provided resources.

